# A chip-based array for high-resolution fluorescence characterization of free-standing horizontal lipid membranes under voltage clamp

**DOI:** 10.1101/2022.04.18.488685

**Authors:** Tobias Ensslen, Jan C. Behrends

## Abstract

Optical techniques, such as fluorescence microscopy, are of great value in characterizing the structural dynamics of membranes and membrane proteins. A particular challenge is to combine high-resolution optical measurements with high-resolution voltage clamp electrical recordings providing direct information on e.g. single ion channel gating and/or membrane capacitance. Here, we report on a novel chip-based array device which facilitates optical access with water or oil-immersion objectives of high numerical aperture to horizontal free-standing lipid membranes while conrolling membrane voltage and recroding currents using micropatterned Ag/AgCl-electrodes. We demonstrate both wide-field and confocal imaging, as well as time-resolved single photon counting on free-standing membranes spanning sub-picoliter cavities are demonstrated while electrical signals, including single channel activity, are simultaneously acquired. This optically addressable microelectrode cavity array will allow combined electrical-optical studies of membranes and membrane proteins to be performed as a routine experiment.

## 1 Introduction

Many membrane processes, including the gating of ion channels, formation of protein pores and membrane fusion give rise to electrical signals such as current flow or membrane potential changes as well as capacitance changes. These signals can be detected with sub-millisecond time resolution and are often directly related to structural changes on the single molecule level. On the other hand, after labelling of either membrane lipids or protein residues with fluorophores, these molecular structural rearrangements can also be followed optically, e.g. using steeply distance-dependent dipole coupling phenomena such as Förster resonance energy transfer (FRET).^1^ There is general consensus that combining electrical and optical readouts of membrane-delimited single-molecule dynamics would be of great value and a number of attempts have been made in this direction which are well summarized in a recent review.^2^ In fact, the combination of single ion channel recording and single molecule fluorescence spectroscopy has been suggested almost 30 years ago.^3^ While patch-clamp fluorometry has already allowed a correlation of protein structural dynamics and functional activity on the ensemble level^4^ and voltage clamping of excised cellular membrane patches routinely yields functional single-molecule resolution for ion channel gating, regularly achieving a recording configuration simultaneously enabling the high optical resolution and sensitivity required for, e.g., single molecule FRET experiments, has proven difficult. Likely for this reason, correlation in membranes between electrical signals reporting on single channel function and optical signals related to structural dynamics has only been shown for a model peptide channel, gramicidin,^5,6^ but not for protein ion channels.

The particular difficulty of experiments combining voltage clamping and optics for single molecule experiments lies in the fact that conditions over which the experimenter has relatively little or almost no control are seldom right for both measurements. E.g. a healthy membrane with high resistance suitable for high-resolution voltage clamping with the right number (often only one) of electrically detectable channels or pores inserted may at long last have been obtained, but the fluorophore on that pore or channel may already have been bleached, or may bleach immediately after recording has really commenced. Conversely, the number of healthy fluorophores may be just right but the membrane has become unstable, leaky or noisy, possibly from photo-oxidation, so as to prevent an actual measurement^2^. Put generally, the product of probabilities that each one of the recording modes yields useful data, which themselves are products of non-unity probabilities, becomes very small and the number of trials to successfully complete a project becomes inordinately large. In order, therefore, to become more routine, such experiments need to be carried out in parallel (array) formats in order to push the success probability into a tolerable range.

A very important requirement for such array formats is, however, that they do not compromise on the resolution of the original methods. As has been pointed out in a recent review^2^ another difficulty in simultaneously applying two highly sensitive and physically distinct measurements to membranes lies in the fact that the requirements for one and the other are not necessarily compatible, especially in miniaturized and parallel formats. For instance, high-resolution voltage-clamp measurements require non-polarizable metal electrodes which are optically non-transparent yet should be as close as possible to the membrane to reduce stray capacitance and access resistance. Efficient collection of photons with high numerical apertures, on the other hand, requires a short working distance on the order of 100 μm which severely limits the space in which electrodes can be arranged. Recently, the droplet-on-hydrogel configuration has been utilized in array formats to perform simultaneous optical and electrical recordings. The prime advantage, here, is that total internal reflection microscopy can be used on these membranes in a straightforward way. However, there are some disadvantages. In order to perform electrical recordings, each droplet has to be manually contacted with an Ag/AgCl electrode. The area of the bilayer is not well controlled and relatively large (>100 μm), decreasing the usable bandwidth for electrical recordings due to high stray capacitance. Not least, formation of the bilayers is a rather intricate process requiring formation of droplets in a separate chamber and considerable waiting time.^7^

Here we show that the architecture of microelectrode-cavity arrays^8,9^ now in routine use for electrical recordings from channels and biological nanopores reconstituted in bilayers^10,11^ can be adapted to provide optical access for high-resolution fluorescence microscopy while maintaning the prerequisites for electrical recording with single-channel resolution.

## 2 Materials and Methods

### Electrophysiology

Voltage-clamp recordings were performed using a modified version of the OrbitMini device (Nanion Technologies, Munich, Germany) equipped with MECA-opto microelectrode cavity arrays (Ionera Technologies, Freiburg, Germany) described below. General experimental procedures can be found elsewhere^10,11^ The 4-channel patch clamp amplifier integrated in the OrbitMini was operated using the Elements Data Reader 3 software (Elements SRL, Cesena, Italy) at 2 nA range with 0-20 kHz bandwidth and 10 kHz digitization rate. Recordings were performed at room temperature (23-25 °C) in 1 or 4 M KCl (pH 7.5, 25 mM TRIS-HCl) or in PBS-buffer as indicated.

The high-bandwidth electrical recording shown in Fig. 4E was performed on a MECA16 microelectrode cavity array (Ionera Technologies). Briefly, one of 16 coplanar gold lines leading to MECA16-cavities (50 μm diam.) was directly contacted with a 1 cm unshielded silver wire to the head-stage of an Axopatch 200B (Molecular Devices, Sunnyvale, CA, USA) patch clamp amplifier operated in capacitive feedback mode at 50 mV/pA gain with the internal low-pass-filter set to 100 kHz cut-off (−3 dB). The output signal was passed through an external low-pass Bessel filter (npi electronic, Tamm, Germany, 8 pole, custom version of LHBF-48X-8HL) set to 50 kHz cut-off frequency and digitized at 1 MHz sampling rate using a PCI-6251 16 bit ADC interface (National Instruments, Austin, TX, USA) controlled by GePulse software (Michael Pusch, University of Genoa, Italy).

### Channel-forming peptides and fluorophores

Native ceratotoxin A (CtxA) was a kind gift from Michael Meyer (Adolph Merkle Institute, Fribourg, Switzerland). Fluorescently labeled CtxA-derivatives were custom synthesized and purified by HPLC (Biosynthan GmbH, Berlin, Germany). Lipid bilayers were formed from di-phytanoyl-phosphatidylcholine (DPhPC) dissolved at 5 mg/ml in octane. Lipid membranes were fluorescently labelled with lissamine-rhodamine-phosphoethanolamine (Liss-Rhod-PE) at <1 % (V/V). Lipids were purchased from Avanti Polar Lipids, Inc (Alabaster, AL, USA), dried as a film and stored at −20 °C under argon atmosphere prior to use.

### Fluorescence microscopy

Epifluorescence wide-field microscopy was performed on an Axiovert200 microscope (Carl Zeiss AG, Jena, Germany) equipped with an Olympus (Tokyo, Japan) PLAPON60XOSC2 (60X/N_a_1.4) oil immersion objective, a HBO100 mercury illuminator (Carl Zeiss AG, Jena, Germany) and a Prime 95B CCD camera (Teledyne Photometrics, Tucson, AZ, USA). Confocal single-photon fluorescence lifetime measurements were performed on a Microtime200 platform (PicoQuant GmbH, Berlin, Germany) using picosecond pulsed excitation at 531 +/- 3 nm, an Olympus UPLSAPO 60X/ N_a_1.2 Ultra-Planachromat water immersion objective, a 50 μm pinhole and single photon detection with Excelitas (Waltham, MA, USA) Avalanche Photodiodes.

### Data analysis and graphing

Wide-field images and stacks were processed in ImageJ (Wayne Rasband, National Institutes of Health, Bethesda, USA). Confocal life-time and photon-counting data were processed in SymPhoTime 64 (Picoquant). Current and voltage traces were exported as axon binary files and imported into IgorPro8 (Wavemetrics, Portland, OR, USA) using Neuromatic XOP (Rothman and Silver 2018). Figures were assembled in PowerPoint (Microsoft, Seattle, USA).

## 3 Results and Discussion

Here, based on the microelectrode cavity array (MECA) described earlier^8,9^ and currently used as a consumable in several commercially available devices for multichannel lipid bilayer electrophysiology^10,11^, we introduce an novel array format that is amenable to high-resolution optical and electrical recording. The MECA principle is based on blind cavities with diameters between 200 and 10 μm in a photolithographically structured 20 μm thick dielectric photoresist (SU8) layer on a 500 μm-thick glass substrate or printed circuit board (PCB). In the original MECA design, the bottom of each cavity consists of a disk-shaped Ag/AgCl microelectrode patterned on a that functions as the active *trans*-side electrode in a voltage-clamp circuit, while a single counter electrode is connected to a common bath on the *cis*-side of the membrane. Bilayers can easily and reproducibly be formed over the cavities using manual and automated variants of the classical Müller-Rudin painting method.^9,12^This configuration with disk-shaped electrodes is amenable to upright microscopy with a water-immersion objective with millimeter working distance and correspondingly low numerical aperture. For high-resolution optical recordings a different approach is required. Here, we present a first solution based on ring-shaped microelectrodes, that define an optical window through glass substrates of coverslip (i.e 170-200 μm) thickness through which the bilayer can be imaged using high NA immersion objectives (**Figure 1**).

**Figure 1.**
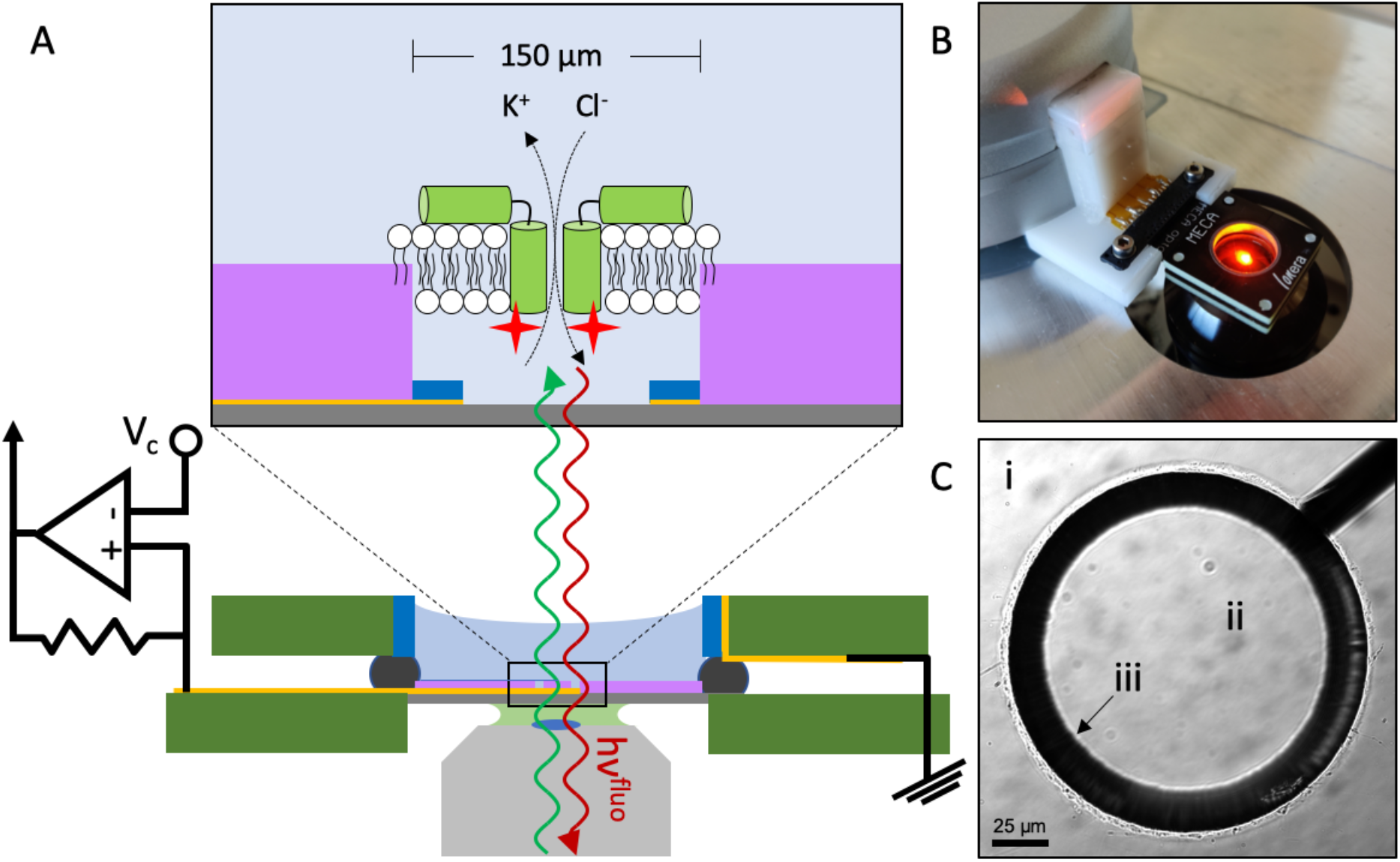
**A:** Schematic (not to scale) of the MECA-opto-device. Excitation and fluorescence light paths in and out of the oil or water (light green) immersion objective (grey) are indicated by green and red waves (hν^exc^, hν^fluo^). Inset: schematic blow-up (not to scale) of one of four MECA-opto cavities. The planar Ag/AgCl microelectrodes (dark blue) and cavities in a 20 μm-thick photoresist (SU8, purple) are structured by lithography, Ag-deposition on Au strip-lines and chloridization as shown previously (Baaken et al. 2008). To allow high-NA optical microscopy, the glass thickness is reduced to <= 200 μm and the electrodes are ring rather than disk-shaped. The devices use in this proof-of-principle demonstration had cavities with diameters of 150 μm. The 12 by 12 mm glass chip is glued into an opening in a fibre-glass/epoxy (FR-4 printed circuit (PCB) material) board on which 5 planar connecting pins are realized. Four of the pins are connected to the gold strip lines leading to the microelectrodes. A rubber O-ring and a circular (10 mm diam.) opening in a second PCB-board define an upper (cis-side) chamber. The inner PCB chamber wall is lined with an Ag/AgCl coating serving as the common ground or bath electrode (dark blue). During assembly, this electrode is connected to the fifth connector pin on the lower PCB-board. B: Photograph of the MECA-optochip mounted into the adaptor mediating connection to a Nanion Orbit-mini recording device placed on the stage of the inverted microscope. The adaptor is constructed so that the lower surface of the MECA.opto-chip is flush with the microscope stage. C: Differential interference contrast (DIC) image taken with an upright microscope of a single MECAopto cavity with SU8 surface (i), glass bottom (ii) and Au/Ag/AgCl ring electrode at the bottom of the cavity (iii). Note that the SU8 surface was put into focus, thus the electrode 20 μm below appears out of focus.

Lipid bilayers can be formed on these apertures by manual “painting” from lipid-in solvent suspensions (Müller-Rudin method) either using a teflon spatula or an air bubble at the tip of a plastic micropipette.^10,11^. As shown in Fig. 2, bilayer formation can be readily ascertained by recording the current response to a sawtooth voltage command resulting in rectangular capacitive current responses from which the size of the thinned-out region of bilayers can be estimated. In the example shown in Fig. 2A, the bilayer was formed on an aperture with diameter 150 μm. The capacitance measured (75 pF) would correspond to a circular bilayer with diameter of approximately 150 μm according to estimates of the specific capacitance of solvent-containing DPhPC-bilayers of approximately 0.4 μF/cm^2^,^13^ while DPhPC bilayers formed by the Müller-Montal technique^14^ have specific capacitances of 0.95 μF/cm^2^ due to lower solvent content^15^. Previous capacitance estimations for DPhPC bilayers on MECAs also indicated that the aperture area is almost entirely occupied by a bimolecular layer and that the area occupied by the solvent annulus is minimal.^16^

**Figure 2.**
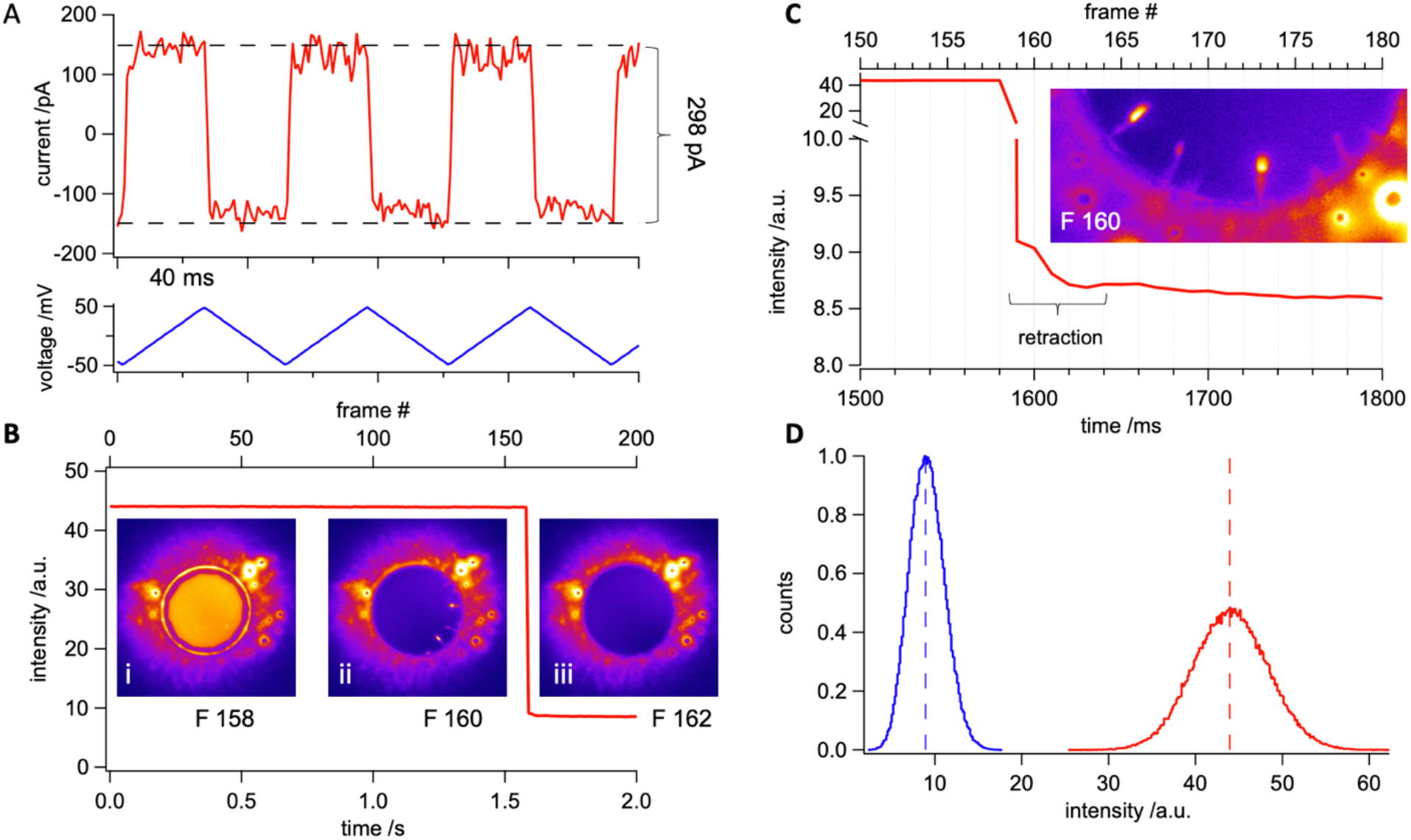
**A:** current vs. time-trace (red) for a DPhPC lipid membrane spiked with Liss-Rhod-PE under electrical stimulation using a triangular voltage protocol of ±50 mV with dV/dt=4 V/s (blue) The corresponding 300 pA capacitive current indicates a total membrane capacitance of 75 pF. **B:** fluorescence intensity versus time trace. Sharp intensity decrease at 1.59 s is caused by irreversible electroporation of the bilayer using a voltage pulse of +1000 mV for 200 ms. Insets: Heat-stained images extracted from a 2 s long sequence (frame rate 1/10 ms) reveal a homogeneously thinned and Liss-Rod-PE spiked DPhPC bilayer (F158) bursting under the voltage. Excess lipid could be observed to retract with finger-like structures towards the SU8 surface lining the aperture (F160) until a fully open orifice with low background signal is detected (F162). **C:** sequence of the fluorescence-time trace immediately preceding and following electroporation to illustrate lipid retraction over 50 ms; inset: rotated blow-up of the rim from frame 160 in B. **D:** fluorescence intensity histograms before (red) and after (blue) membrane electroporation. A clear shift towards lower greyscale values indicate loss of the fluorescence signal.

Inset i in Fig. 2B shows a wide-field fluorescence micrograph from the same membrane that provided the electrical recording in Fig. 2A and that was labelled by admixing 0.5% lissamine-rhodamine phosphoethanolamine to the lipid suspension used for membrane formation. In order to reduce background, the field of illumination was limited to the center of the cavity using an iris. A sharp projection of the iris in the plane of the membrane was also helpful in setting the focus. The graph in Fig 2B shows the fluorescence intensity in that area plotted against time and frame number (frame rate 100 Hz). At frame 159 = 1.59 s the membrane was destroyed by electroporation with a pulse of high voltage (1 V). resulting in a 80% loss of fluorescence (see Fig. 2D). Interestingly, at the frame rate used, an intermediate stage can be observed (Fig. 2Binset ii, Fig. 2C) where fluorescent lipid can be seen to retract in radial fashion towards the rim of the aperture. Complete retraction of lipid took place within 5 frames (159-164) corresponding to the surprisingly long interval of 50 ms.

The ability to simultaneously observe optical and electrical dynamics in free-standing membranes is particularly attractive for studying molecular events leading to the opening of voltage-dependent ion channels. In particular, correlating the movement of single voltage sensor domains detected optically with open-closed transitions measured electrically would be a major step forward. Antibiotic polypeptides such as mellitin^17^ and alamethicin^18–20^ form multimeric channels in a voltage-dependent manner and can be regarded as models for voltage-dependent gating. Both fluorescently labelled alamethicin^21^ as well as mellitin^22^ have been used in fluorometric measurements on free standing lipid membranes; in the case of mellitin, channel activity was successfully correlated on the ensemble level with optical detection of voltage-dependent peptide reorientation. For single channel studies, melittin is not ideally suited because the current steps are not as clearly delineated as those for alamethicin. On the other hand, the chemical nature of alamethicin as a peptaibol containing non-amino-acid monomers complicates synthesis of fluorescent derivatives. Recently, however, a further voltage-dependent antibacterial peptide, Ceratotoxin-A (CtxA), –natively found in the mediterranean fruit fly *Ceratitis capitata*–was described that consists entirely of proteinogenetic amino acids but shows well-defined current levels like alamethicin.^23–25^

In order to ascertain the possibility of performing correlated single-channel and fluorescence recording, we therefore prolonged the native 36 amino acid sequence of Ctx-A was prolonged by a C-terminal cysteine, which was then fluorescently labeled with an Atto532-maleimide dye at its thiol (CtxA-C-Atto532M). Based on structural and functional similarities to alamethicin, CtxA is conceptualized as consisting of two helices, of which the C-terminal one, featuring mainly hydrophobic residues, might reorient into the membrane, while the N-terminal helix, possessing multiple charged lysine residues, remains on top of the membrane. According to the current model, which is analogous to the barrel-stave model for alamethicin channel formation, a minimum of three CtxA monomers, partly reoriented into the membrane, will then form a transient pore with a central channel, which can be discretely increased in its diameter due to random incorporation of further reoriented monomers.^23–25^

**Fig. 3A** shows a representative 3 s current-time trace recorded from a DPhPC-bilayer in the presence of 5 nM CtxA-C-Atto532M revealing burst-like current transients due to CtxA assembly and pore opening at a holding potential of +140 mV *trans*. At the end of the recording, the membrane was again irreversibly electroporated which resulted in a sharp current increase (see also **Fig. 3B**) as well as an abrupt decrease in the fluorescence signal (**Fig. 3C**).However, as is also evident from the fluorescence images before and after membrane disruption (**Fig. 3 D**) the relative reduction in brightness was much less than that observed with fluorescent lipids.. As revealed by confocal microscopy (see below), the autofluorescence of borosilicate glass probably contributed considerable background.

**Figure 3.**
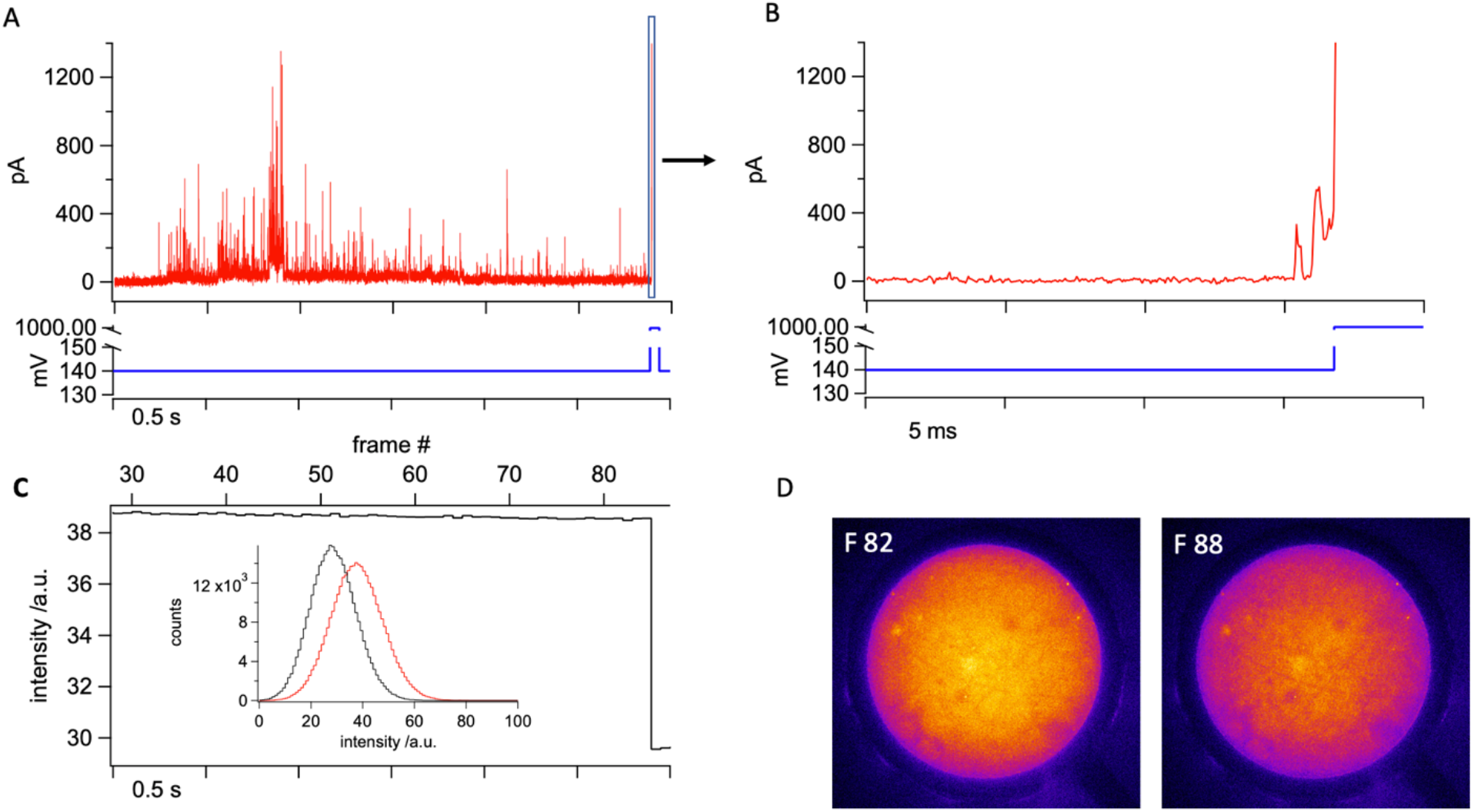
**A:** current-time trace (red) recorded from a DPhPC bilayer at a bandwidth of 10 kHz and 20 kHz sampling rate in 1 M KCl incubated with 5 nM CtxA-C-Atto532M under a constant trans-positive voltage of +140 mV trans (blue). Transient current events are caused by spontaneous pore formation of membrane associated CtxA peptides under the applied voltage. The bilayer was sacrificed by the end of the experiment using a 200 ms long voltage pulse of 1000 mV; **B:** expanded current and voltage traces during from the region indicated by the black box contour in panel A. Abrupt current increase indicates loss of the bilayer. Prior current peaks are caused by CtxA pore formation; **C:** fluorescence over time trace extracted from a 6 s long movie sequence during burst of the bilayer. The sudden drop of fluorescence intensity indicates the loss of the bilayer. Inset shows histograms of fluorescence intensity in the membrane area before (red) and after (blue) membrane loss. **D**: Two images from the sequence obtained before (82) and after (88) membrane destruction.

In order to further analyze the relative contribution of membrane-bound channel-forming peptide to the fluorescence signal, we conducted a single-photon counting fluorescence life-time study on free standing bilayers, comparing DPhPC membranes either spiked with Liss-Rhod-PE or doped with CtxA-C-Atto532M. Fig. 4A shows a representative fluorescence lifetime image (FLIM) recorded in the x-z plane from a free standing Liss-Rhod-PE spiked DPhPC membrane. The scan is situated at the border of the cavity, and shows the SU8-layer to the right displaying, as expected, considerable fluorescence with excitation at 530 nm^26^ as does the glass bottom of the cavity. The membrane (M) can be seen to separate the *trans*- from the *cis*- volume; its fluorescence is very weak above the electrode (E) due to the shadow cast by the metal. Interestingly, the membrane appears curved outward. Some degree of membrane curvature, either inward or outward (see Fig. 5 A) was observed repeatedly in these experiments. Repeated formation of membranes on the same aperture could lead to inwardly and outwardly curved as well as flat membranes, suggesting that the precise movement by which the membrane is manually formed influences its geometry. Indeed, a similar observation was made in a previous confocal study on freestanding lipid bilayer membranes.^27^ Importantly, the *cis* and *trans* volumes above and below the membranes were free of detectable fluorescence. It is also obvious that the membrane appears surprisingly thick with a full width at half maximal intensity of 3.7 μm compared to membranes from e.g. giant unilamellar vesicles.^28^ This finding, which is in line with previous results^27^ is probably partly due to the inherently inferior resolution in the z-axis compared to that of the x-y plane in confocal measurements, as well as to relatively coarse scanning step used (300 nm). Labelled lipid is also clearly visible on top of the SU8 layer (compare Fig. 2B and Fig. 5 in Ref. which has the expected thickness of approximately 20 μm. Labelled lipid is seen to be present on the SU8 surface (compare Fig. 2B) as well as at the rim of the aperture, likely corresponding to the torus or annulus.^29^ The borosilicate glass layer can be clearly seen to emit fluorescence with lifetimes considerably lower than that of the membrane.

**Figure 4.**
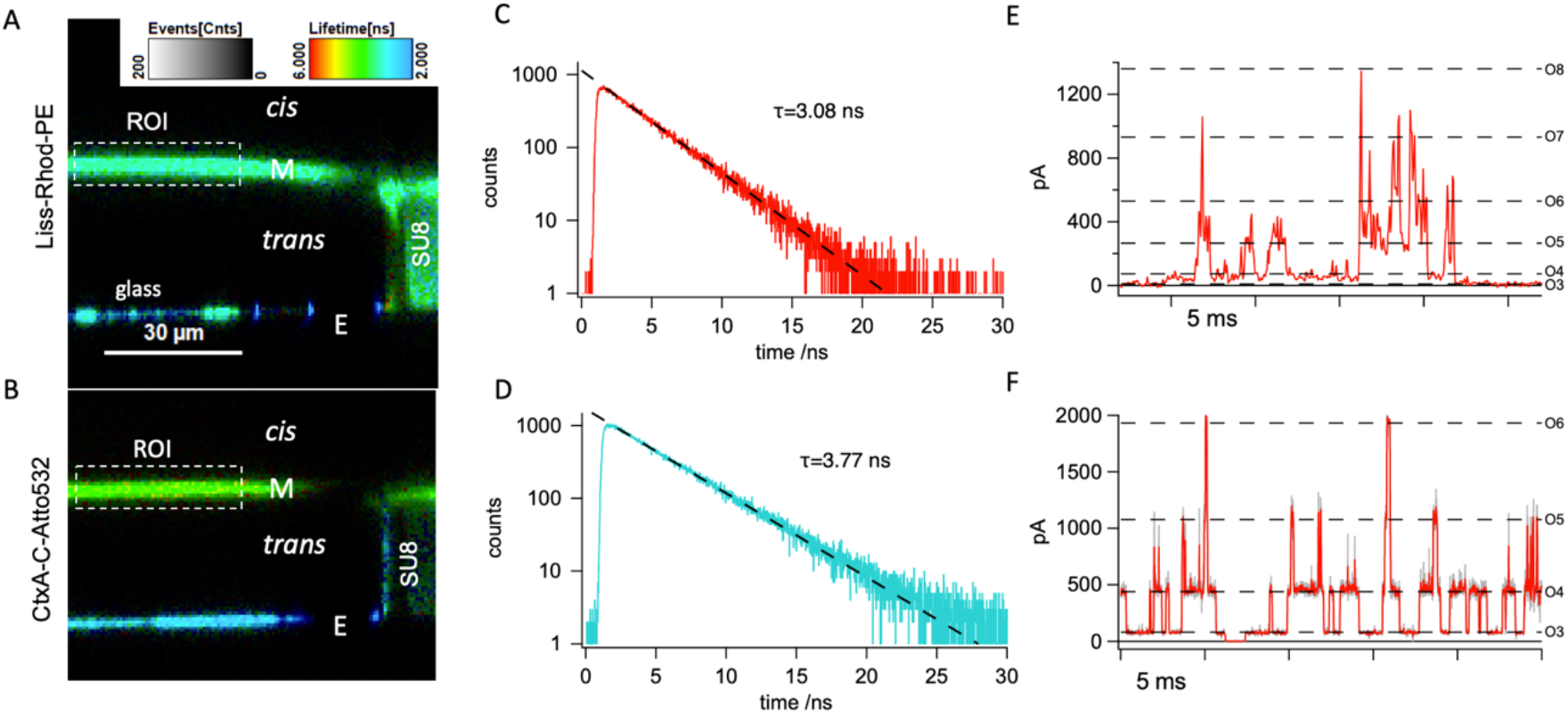
**A:** confocal x-z fluorescence-lifetime-scan of a freestanding DPhPC membrane spiked with 0.5% lissamine-rhodamine-phophatidylethanolamine (Liss-Rhod-PE) spanning a 150 μm SU8 (turquoise, right side) aperture over a glass chip (blue, bottom). The image was acquired in PBS buffer at pH 7.5; **B:** confocal x-z fluorescence-lifetime-image of a freestanding unstained DPhPC membrane intoxicated with 5 nM CtxA-C-Atto532M (green). Notably, the electrode induced shading had a major effect on the fluorescence at the SU8-lipid interface due to the fact, that the dye was distanced from the SU8 surface by the lipid, thereby reducing unspecific reflections in this area. The clear signal of short lifetimes (blue) indicated the glass substrate. The image was acquired in 1 M KCl at pH 7.5; **C:** fluorescence lifetime histogram obtained from the membrane region of interest (ROI) shown in A in a single photon counting experiment. Exponential fitting revealed a lifetime of 3.08 ns. D: fluorescence lifetime histogram obtained from the membrane region of interest (ROI) shown in B. Exponential fitting revealed a lifetime of 3.77 ns. E: representative current-time trace recorded during simultaneous optical membrane investigation in 1 M KCl at a constant trans-positive voltage of 120 mV showing transient pores are formed by fluorescently labeled CtxA-C-Atto532MCurrent recordings were carried out with the Orbitmini device at a final bandwidth of 20/2=10 kHz; **F:** exemplary current over time trace recorded with native CtxA in 4 M KCl from a DPhPC bilayer immersed in 4 M KCl using a trans-positive voltage of 140 mV at a sampling-rate of 1 MHz, low-pass filtered at f_c_(−3 dB)=50 kHz (grey) and 25 kHz (red). In E and F, current levels open states corresponding to trimer (O3) to octamer (O8) (dashed lines) were derived by linear correction for the experimental conditions used here of values reported in Ref.^23^.

**Fig. 5.**
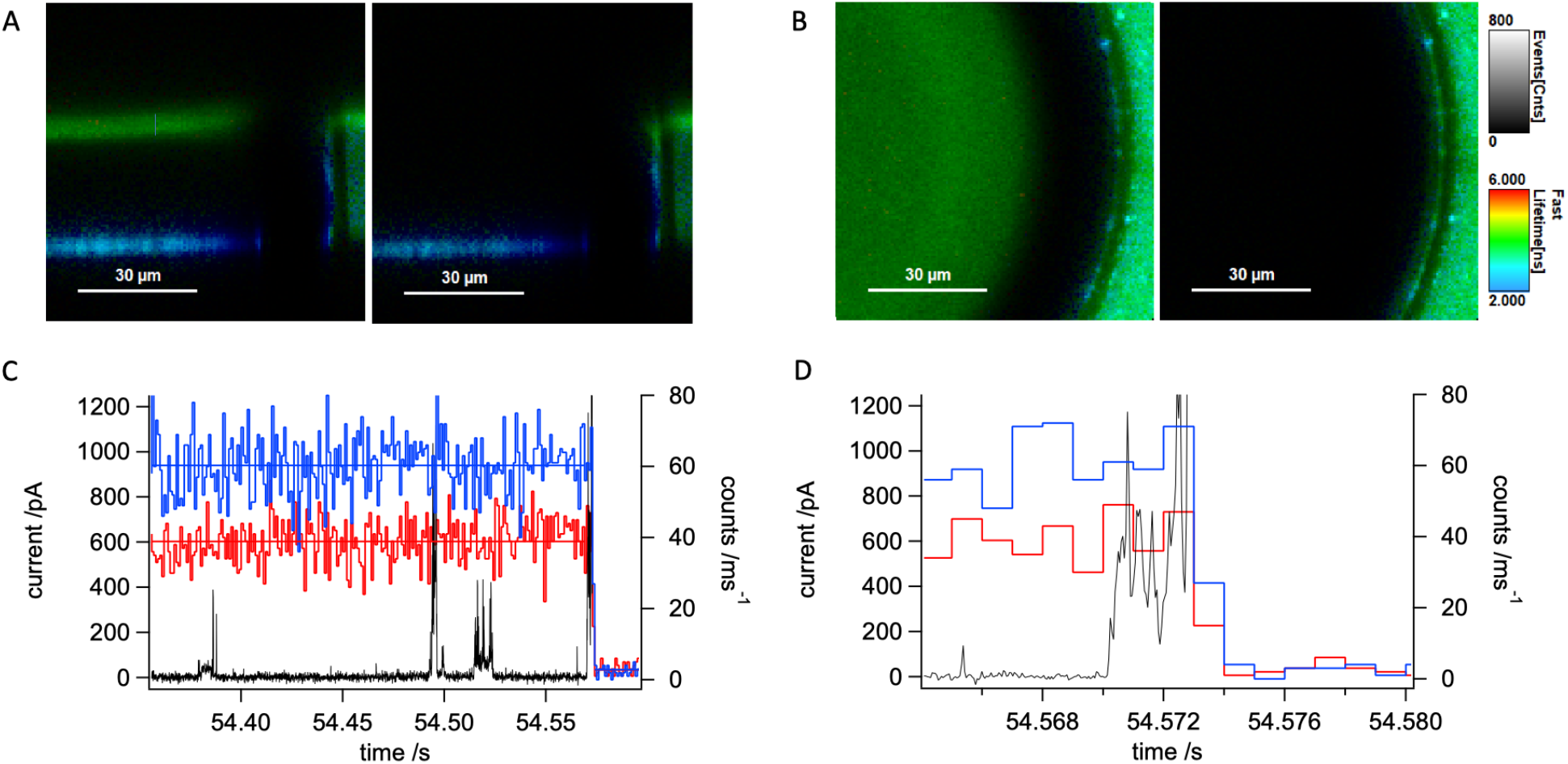
**A,B**: confocal x-z (A) and x-y(B) fluorescence lifetime imaging scan of a microelectrode cavity with membrane (left panels) and after spontananeous membrane rupture (right panels) in the presence of 5 nM CtxA-C-Atto532M in the cis-compartment. C, D: photon count (blue: vertical polarization, red: horizontal polarization) and ion current-time traces simultaneously recorded from the membrane shown in A,B.

Fig. 4b shows another cavity containing a pure DPhPC-bilayer after equilibriation with 5 nM CtxA-C-Atto532M added to the *cis*-compartment. Fluorescence can clearly be detected in the membrane as well as above the SU8-layer indicating enrichment of fluorescent peptide at the lipid/water interface. The annulus is not visible, as expected when fluorescent material is present only at the *cis*-ward surface. Again, and in line with the wide-field epifluorescence results, there is considerable fluorescence of the glass bottom.

Single-photon counting lifetime measurements in the region of interest revealed a characteristic lifetime of approximately τ=3 ns for fluorescence emitted from Liss-Rhod-PE spiked DPhPC membranes (Fig. 4C), which is in good agreement with the literature value for the conjugated fluorophore.^30^ The lifetime of the fluorescence observed in CtxA-C-Atto532M-doped membranes (Fig. 4D) was τ=3.8 ns in perfect agreement with the value given by the manufacturer for the free Atto532 dye.

Current measurement from the membrane shown in B at a holding potential of +120 mV *trans* revealed discrete current levels, characteristic for CtxA (Fig. 4E). Comparison with native CtxA recordings shown in Fig. 4F revealed similar behavior confirms the overall function of the labeled CtxA to be unaltered. Note that the recordings of native CtxA were performed at a bandwidth of 50 kHz so that we were able to fully resolve the sub-ms visits to open levels >O3, which was not possible with the lower bandwidth of the recordings with the Orbit mini (Fig. 4E). This is due mainly to the fact that the first generation of MECA-opto devices used here has apertures of 150 μm diameter, i.e. three time that of the MECA16 used, and that the resulting quadratic increase in capacitance limits the usable bandwidth due to increased noise.^31^

While CtxA-C-Atto532M was able to produce channel openings compatible with the native peptide, we noted a heightened tendency of the membrane to break spontaneously in the course of channel activity suggesting that either the cysteine or the fluorophore interacts with and destabilizes the membrane upon channel formation. Fig. 5 shows a further experiment using CtxA-C-Atto532M. The membrane, imaged in x-z and x-y in the left panels of Fig. 5A and B, respectively, is seen to slightly curve into the cavity. Fig 5C and D show traces of photon count (kHz) obtained with two avalanche photodetectors at vertical (blue) and horizontal (red) polarization as well as the simultaneously acquired current signal at a constant voltage of +120 mV cis. Note that we were unable to observe any changes in fluorescence intensity upon changing the voltage between −120 and +120 mV. Before the membrane breaks the photon counts remain stable at 60.31 kHz (vertical polarization) and 38.89 kHz (horizontal polarization). When the membrane ruptures during the final burst of channel activity at 54.573 s (see also left panels in Fig. 5A,B) both photon counts are strongly reduced to values of 2.82 (vertical polarization) and 2.76 (horizontal polarization). With vertically polarized excitation, there is, therefore, considerable polarization anisotropy (0.155) of the emission from the membrane bound label that is reduced to near 0 (0.007) for emission from the solution (after membrane rupture). This result is compatible with a relatively strong interaction between the fluorophore and the membrane. However, because of the flexibility of the linker between the peptide and the fluorophore (see Fig. SI3), we suspect that the limited mobility of the fluorophore is due to direct interaction between the lipid’s choline and the fluorophore’s sulfite groups rather than mediated by the peptide.

The experiment shown in Fig. 5 also clearly illustrates the fact that at the concentrations necessary to observe ion channel formation, photon counts are too high to permit single molecule measurements such as fluorescence correlation spectroscopy. Similar results were obtained in an early study on fluorescently labelled alamethicin, where only a very small fraction of membrane-associated peptides appear to be involved in channel formation.^21^ It should be noted, however, that this problem is unique to peptides forming multimeric pores and is will not arise with classical ion channels.

## 4 Conclusion

We shave shown fluorescence characterization of free-standing horizontal lipid membranes under voltage clamp by epifluorescence wide-field, as well as time-resolved confocal microscopy with simultaneous electrophysiological measurements using a novel, optically and electrically addressable micro-electrode-cavity-array (MECA-opto). The optical quality of the experimental apparatus introduced here allow confocal fluorescence lifetime imaging, determination of fluorescence lifetimes as well as fluorescence anisotropy from single-photon counting. Wide-field fluorescent microscopy is somewhat disturbed by the autofluorescence of the borosilicate glass bottom, which may be reduced by using quartz glass as a low-autofluorescence substrate.^32^ In addition, the device can easily be produced with smaller apertures to enhance the bandwidth of electrical recording by reducing bilayer capacitance if the necessity arises for very short events to be resolved. However it should be noted that smaller bilayers also reduce the efficiency of ion channel reconstitution, a step that is going to be crucial if simultaneous optical electrical measurements on single protein channels will be attempted. In summary, this device has the potential to greatly simplify simultaneous optical and electrical recordings from ion channels, pore forming peptides and biological nanopores in membranes and to realize a long-standing ambition to correlate structural and functional dynamics of single membrane proteins on the single molecule level.

## Acknowledgements

TE was partly funded by a PhD fellowship in the framework of the International Graduate College 1642 “Soft Matter Science: Concepts for the Design of Functional Materials” of the Deutsche Forschungsgemeinschaft (DFG). Work in JCB’s laboratory was funded by the German Ministry for Research and Education through Project Management PTJ (TseNareo), by the BW Foundation through Project Management VDI (MSDS-BioMem) and by the Ministry of Commerce of the State of Baden-Württemberg in the Framework of the Forum Gesundheitsstandort Baden-Württemberg (TechPatNano). Acquisition of the Microtime 200 was co-funded by a Major Research Instrumentation Grant from the DFG (project No. 290424854) and by the BW-foundation (project BITS). We thank Dr. Gerhard Baaken, Dr. Sönke Petersen and Dr. Ekaterina Zaitseva of Ionera Technologies GmbH for realizing the chip structures and for helpful suggestions.

## Author contributions

T.E. and J.C.B. conceived and designed experimental work, T.E. performed experiments and analyzed experimental data, T.E. and J.C.B. prepared figures and wrote the manuscript.

## Competing interests

J.C.B. is co-founder and shareholder of Nanion Technologies GmbH, Munich, Germany and Ionera Technologies GmbH, Freiburg, Germany

